# A guide to D-SPIN: constructing regulatory network models from single-cell RNA-seq perturbation data

**DOI:** 10.1101/2025.07.28.667326

**Authors:** Jialong Jiang, Matt Thomson

**Affiliations:** Division of Biology and Biological Engineering, California Institute of Technology, Pasadena, CA, 91125, USA; Center for Studies in Physics and Biology, The Rockefeller University, New York, NY, 10065, USA

**Keywords:** D-SPIN, gene regulatory networks, single-cell perturbation experiments, spin networks, Markov random fields, probabilistic graphical models, Perturb-seq

## Abstract

D-SPIN (Dimension-scalable Single-cell Perturbation Integration Network) is a computational framework for constructing regulatory network models from large-scale single-cell perturbation data. D-SPIN models the transcriptional state of cells as a set of individual genes or gene programs that interact through a condition-dependent spin network model or Markov random field. A single unified network is learned across multiple conditions, where the perturbations modulate the activity of individual network nodes. The architecture allows information integration between different perturbation conditions and supports efficient inference algorithms that scale to thousands of genes, thousands of conditions, and millions of single cells. For interpretability, D-SPIN can also operate at the level of programs of co-expressing gene sets, which are extracted via unsupervised orthogonal non-negative matrix factorization (oNMF). Here we present the theory and detailed procedures for applying D-SPIN to build program-level or gene-level network models from single-cell data. With a public immune dictionary dataset, we illustrate multiple applications of D-SPIN, including identifying gene programs and network modules, classifying perturbation responses of cytokines, and nominating key regulators that mediate cytokine responses.

## 1 Introduction

Gene regulatory networks (GRNs) orchestrate gene expression in response to internal and external cues to achieve the physiological functions and survival of cells [1, 2]. However, the organizing principles of GRNs remain poorly understood, especially the global control logic that operates at the transcriptome scale. Recent single-cell RNA-seq approaches with large-scale multiplexed perturbation experiments, such as MULTI-seq and Perturb-seq, enable profiling of the transcriptomic cellular response to thousands of genetic or chemical perturbations [3–6]. Although perturbations are widely applied in genetics and known to be informative in revealing regulatory interactions [7, 8], converting these massive perturbation datasets into cell-scale models of regulatory networks remains a major theoretical and computational challenge.

To address these limitations, we have developed D-SPIN (Dimension-scalable Single-cell Perturbation Integration Network), a novel theoretical framework that integrates multi-condition single-cell data into unified regulatory network models of perturbation responses [9]. D-SPIN models the cell as a collection of interacting genes or gene expression programs and constructs a probabilistic model, known as a spin network or Markov random field, to infer regulatory interactions between genes and applied perturbations. Spin network models originated in statistical physics and have been generalized to many systems and applications via maximum entropy methods in physics and via Hopfield associative memory models in machine learning [10–12]. Spin network models have shown promise in inferring gene regulatory network models from bulk gene expression data using the Hopfield memory storage algorithms [13, 14]. However, the application of spin network models in the single-cell context remains limited due to the lack of an effective statistical inference framework that can scale to large networks, incorporate perturbations, and capture the information contained in the entire cell state distribution measured in single-cell experiments.

To address these challenges, D-SPIN exploits a natural factorization within the mathematical structure of the Markov random field to enable information integration in large datasets with thousands of conditions and millions of single cells. The model learning problem is divided into two parts: (1) construction of a unified regulatory network model and (2) inference of how each perturbation interacts with the network model. In practice, the gradient function of the inference problem is independently estimated on each condition, similar to the minibatch training schemes in deep network architectures. Therefore, training can be parallelized across hundreds of CPU cores, significantly speeding up computation. Depending on the size of the network, D-SPIN employs three training algorithms with increasing scalability: exact maximum likelihood, Markov chain Monte Carlo (MCMC) maximum likelihood, and pseudolikelihood. Each of the methods is guaranteed to converge to the optimal solution with sufficient training data and computation time [15–17]. In benchmarking tests, D-SPIN achieved state-of-the-art accuracy on both synthetic and biological perturbation datasets with hundreds of elements [9, 18]. D-SPIN can also incorporate prior information on network topology during network inference, such as transcription factor (TF) binding motifs or chromatin accessibility data from ATAC-seq, to further improve inference accuracy.

Although D-SPIN scales well to large networks, interpreting network models with thousands of nodes is challenging. We therefore designed D-SPIN to operate on both single genes and gene expression programs. Gene programs are a group of coexpressing genes across perturbation conditions [19, 20]. The program-level description enables D-SPIN to derive a coarse-grained description of a cellular regulatory network in terms of physiological functions and pathways, and their interactions with applied perturbations. Building upon existing algorithms, D-SPIN includes an automatic procedure for identifying gene programs from single-cell data with unsupervised orthogonal non-negative matrix factorization (oNMF) and phenotypic gene-program annotation, and can also accommodate pre-defined gene sets from prior biological knowledge [21–23]. Furthermore, to investigate the mechanisms of program-level response, we developed a computational framework to identify key regulators from the gene-level networks that mediate the observed program responses.

D-SPIN provides an interpretable, generative network inference framework for obtaining insights into the global regulatory mechanisms and identifying how perturbations act on the network. D-SPIN can accommodate a wide range of different perturbation types, including genetic perturbations, small molecules, extracellular signals, and even physiological states of health and disease. In this chapter, we present the theoretical derivation and detailed protocols for using D-SPIN in three scenarios with our public Python package, including (1) constructing program-level network models, (2) constructing gene-level network models, and (3) associating program-level responses with their gene regulators. As a case study, we apply D-SPIN to a public immune dictionary dataset of *in vivo* cytokine responses [24]. D-SPIN reveals the cell-type-based network partitioning of immune response gene programs on CD4, CD8, and regulatory T cells, dissects selected cytokines into three major categories (Th1-polarizing, IL-1 subfamily, and type I interferon), and proposes Stat2-Irf7/Stat1-Irf1 as the key diverging effectors that mediate type I versus type II interferon responses.

## 2 Materials

### 2.1 Software and code

D-SPIN is primarily implemented in Python and released through PyPI and GitHub (github.com/JialongJiang/DSPIN). D-SPIN can be installed through pip with `pip install dspin.` D-SPIN has relatively few dependencies, primarily `scanpy` and other basic scientific computing packages. Nonetheless, we recommend setting up a fresh `conda` or `mamba` environment to avoid version conflicts between packages.

The Python implementation of D-SPIN is sufficient to handle datasets of 100,000 cells in a few hours on a local desktop. For large datasets with millions of cells, we also provide a MATLAB implementation supporting multi-core parallelization to speed up the computation. Using MATLAB’s `parfor` parallelization, users can distribute computations across local CPU cores or cluster nodes. This often yields near-linear speedup with the number of cores utilized.

### 2.2 Input data

D-SPIN is flexible with perturbation types; the main assumption is that all conditions share a common core regulatory network, even if the cell state distributions vary. Examples of perturbations include Perturb-seq, drug treatments, extracellular signals, healthy versus disease conditions, different patients, different time points, or different local microenvironments from spatial transcriptomics. D-SPIN requires single-cell data, as D-SPIN explicitly models the complete distribution of transcriptional states. Performance on bulk sequencing data is more limited due to the loss of distribution information.

The single-cell data should be stored in `AnnData` format with the following attributes

– `adata.obs[‘if_control’]: adata.X`: Single-cell count matrix. Basic preprocessing and quality control steps are assumed, such as filtering for low-quality cells and highly variable gene selection. The count matrix should be normalized by library size and log-transformed.
– `adata.obs[‘sample_id’]` : Label for experimental conditions. D-SPIN is tolerant of conditions with low cell numbers, as the network learning procedure shares information between conditions. Nonetheless, we recommend at least 25 cells per condition..
– `adata.obs[‘batch’]` : Label for experimental batches. A batch refers to cells processed together in the same experimental run. Perturbation responses are primarily compared within the same batch for batch effect correction.
– `adata.obs[‘if_control’]:`Label for whether the experimental condition is a control. Perturbation responses are compared with the control samples to obtain the relative response of perturbations. If no explicit control sample is available, perturbations are compared with the average of all conditions.

## 3 Methods

### 3.1 Overview of D-SPIN

#### Theory of the D-SPIN framework

Here, we develop the mathematical framework and computational strategies to construct a gene regulatory network model from many single-cell perturbation conditions. Mathematically, D-SPIN builds a spin network model or Markov random field, which can be interpreted as an interaction network. Spin-network models were initially introduced to study magnetic materials known as spin glasses, whose properties emerge through positive and negative interactions between localized magnetic moments or spins [15]. Spin network models have been generalized and applied to many systems of interacting elements in physics and machine learning, including neural networks, bird flocks, and proteins [12, 14, 25], and also to study the storage of memories known as Hopfield networks [10]. However, the application of spin network models to single-cell data has been limited by the mathematical and computational challenges of solving the inference problem on large datasets.

In general, spin network models consider a network of *M* interacting elements *s*_*i*_. The goal of the framework is to assign a probability *P* (*s*_1_, …, *s*_*M*_) to each configuration of the network with an energy function. The energy function *E*(***s***) computes the effective energy of a given transcriptional state ***s*** by balancing the influence of the interactions between elements (encoded in ***J***) and the activity of individual elements (encoded in ***h***). The resulting probability of a given transcriptional state ***s*** is

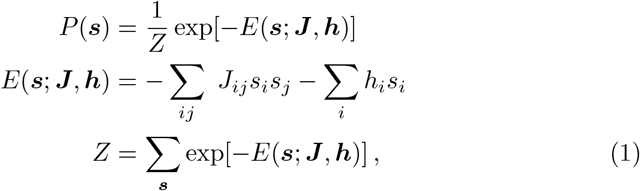

where *Z* is a normalizing constant ensuring all probabilities sum to 1. The regulatory network ***J*** accounts for activating (*J*_*ij*_ > 0) and inhibitory (*J*_*ij*_ < 0) interactions between network nodes. The vector of activity ***h*** encodes a bias on each element: a positive *h*_*i*_ increases the likelihood of “on” state (high expression, *s*_*i*_ = 1), whereas a negative *h*_*i*_ promotes “off” state (low expression, *s*_*i*_ = −1 (Fig. 1A).

**Fig. 1.**
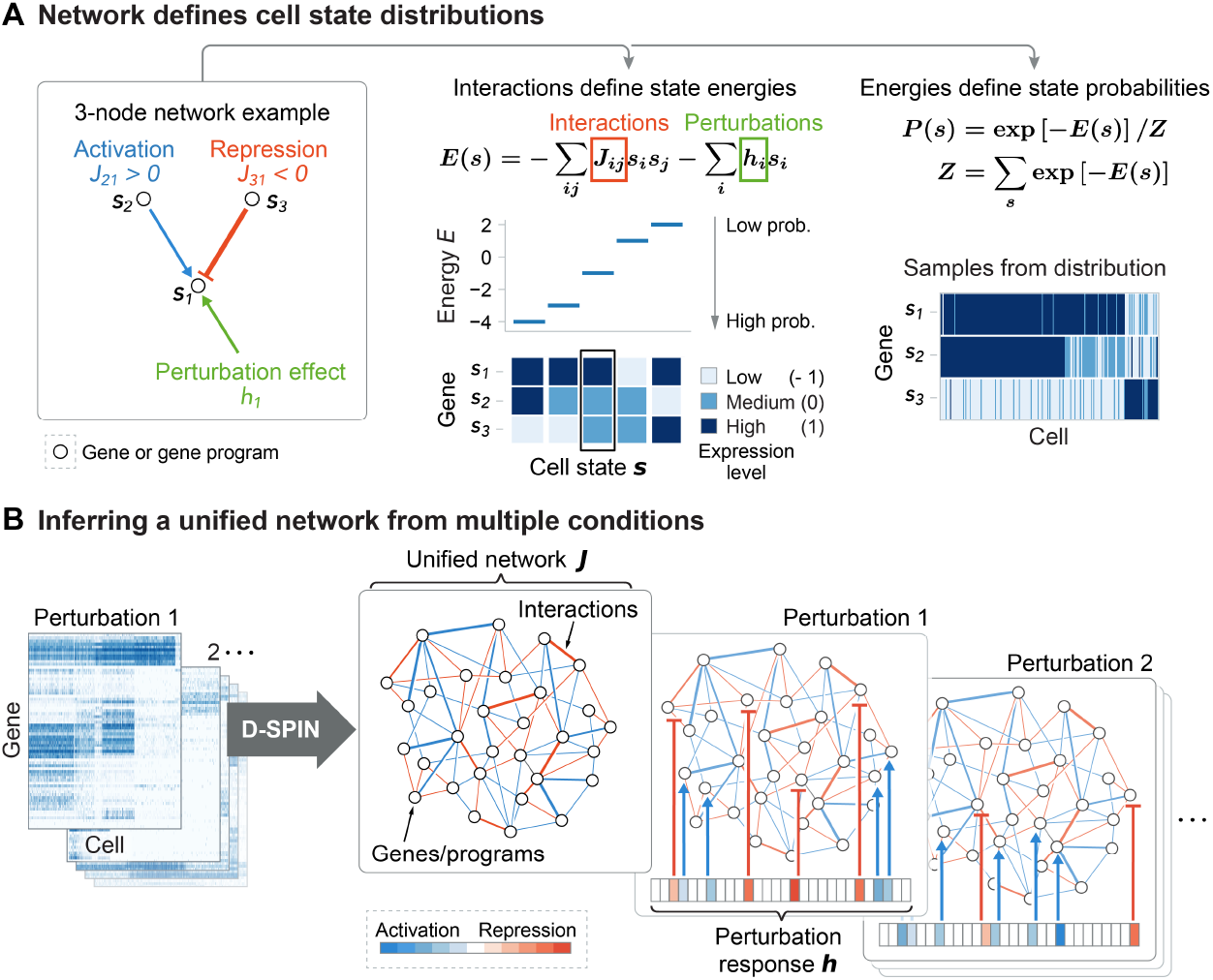
Schematics of the D-SPIN framework. (A) The interactions between genes or programs and the perturbation effects define an energy function *E*. Cell states with lower energy have higher probabilities. The energies together define a distribution over all states. For instance, in the 3-node network *J*_21_ = 1, *J*_31_ = 2, *h*_1_ = −1, the energy of the boxed cell state ***s*** = [1, 0, 0]^T^ is *E* = −*J*_21_*s*_1_*s*_2_ − *J*_31_*s*_1_*s*_3_ − *h*_1_*s*_1_ = −1. (B) D-SPIN infers a unified network model across perturbation conditions. The network ***J*** remains constant while the activities of individual genes or programs are modulated by the perturbation ***h***.

The key insight in D-SPIN is that the regulatory network remains largely conserved across conditions, while each perturbation modulates the activities of individual genes to activate different internal modes of the network, creating altered transcriptional state distributions. Specifically, all conditions share a unified ***J*** while each perturbation *n* has a distinct ***h***^(*n*)^ (*see* **Note 1**) (Fig. 1B). Conceptually, the interaction matrix ***J*** defines an energy landscape of possible transcriptional states, while the perturbation effects ***h***^(*n*)^ apply linear tilts to the landscape, biasing the cell state distributions. Specifically, D-SPIN optimizes the following joint log-likelihood function

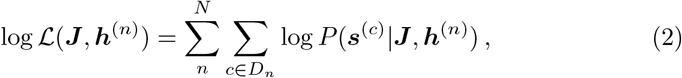

where *c* is cell index and *D*_*n*_ is the *n*-th experimental condition.

In D-SPIN, *s*_*i*_ are chosen to be three-state discrete variables as [−1, 0, 1] instead of two-state variables as [−1, 1] common in statistical physics. The three-state representation captures intermediate expression levels, which can be functionally relevant for gene regulation, and it also improves quantitative agreement between model and experimental cell state distributions (*see* **Note 2**). Similar choices of discretization have been deployed to reduce noise and accommodate nonlinear dependencies from the pioneering works in modeling gene expression networks using microarray measurements [26]. In practice, we discretize the continuous expression level of genes or programs into three states with K-means clustering (*see* **Note 3**).

#### Inference algorithms for D-SPIN

The mathematical form of D-SPIN not only provides a natural and interpretable representation of perturbation action, but also enables us to develop efficient and scalable learning algorithms by separating the training of each condition. Training of general graphical models typically requires estimating the gradient function for every data point separately, which is computationally expensive. In contrast, the gradient of the D-SPIN objective function is pooled for each experimental condition. Consequently, the training of the network can be deployed in parallel, with each computational thread computing the gradient for a single experimental condition. The approach eliminates the need to estimate gradients on each single cell separately, and minimizes data communication cost by only passing the computed gradient of the compact number of parameters (***J*** and ***h***) in D-SPIN. With the parallelization toolbox in MATLAB, the pipeline can easily utilize hundreds of CPU cores, enabling efficient inference in datasets with millions of single cells.

The multi-condition spin network optimization solved by D-SPIN is a convex problem, ensuring that the only local optimum is the global optimum [9]. Therefore, gradient-based optimizers can be used without the risk of getting trapped in local optima. However, challenges remain due to indistinguishable model and floppy model directions [27, 28]. Perturbations that generate diverse transcriptional states can help resolve these issues, leading to more accurate inference results, which is also an important theoretical motivation for D-SPIN (*see* **Note 4**).

Specifically, we adapted three inference methods for D-SPIN, which are selected from the most accurate approaches in the spin network inference literature [15]. Each method has advantages under specific scenarios depending on network size, number of cells, and number of perturbation conditions.

##### Exact maximum likelihood inference

The gradient of the objective function is

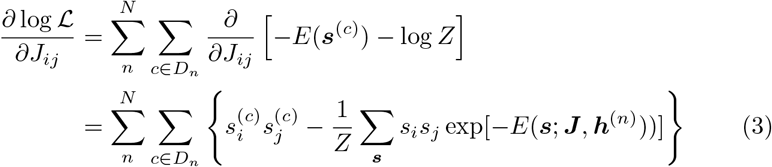

The first term is the sum of *s*_*i*_*s*_*j*_ in all experimental samples, and the second term is a constant, which is the expectation of *s*_*i*_*s*_*j*_ of the current model, defined by the parameters ***J***, ***h***^(*n*)^. We normalize the gradient by cell number and sample number to improve numerical stability under a given step size. With a similar derivation for ∂ log *ℒ/*∂*h*_*i*_, the gradients of the objective function can be written as follows.

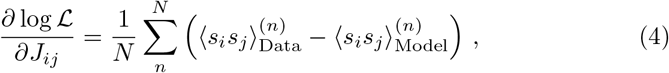

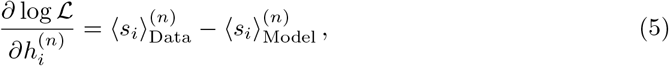

where 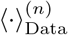is the average over the data of *n*-th conditions, and 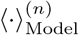 is the expectation on the distribution defined by current model parameters ***J***, ***h***^(*n*)^. Therefore, the modeling learning is essentially matching the cross-correlation ⟨*s*_*i*_*s*_*j*_ ⟩and mean ⟨*s*_*i*_⟩ between the model and data.

For small networks, the probability distribution *P* (***s J***, |***h***) can be explicitly computed, thus the exact gradient of ***J***, ***h*** can be computed using Eq. 4, Eq. 5. Once the gradients are computed, any standard optimizer such as gradient ascent, Momentum, or Adam can be deployed to solve the optimization problem. As the gradient estimation requires enumerating all possible states, the computational complexity scales exponentially with the number of nodes. In practice, the method is only applicable to small networks of less than 15 nodes (3^15^ ≈1.4 × 10^7^ states).

##### Markov Chain Monte Carlo (MCMC) maximum likelihood inference

As the gradient only requires computing the mean and cross-correlation of the samples, the complete distribution *P* (***s|J***, ***h***) can be approximated by drawing samples from an empirical distribution. Without evaluating the exact distribution, we can construct a Markov Chain over states whose stationary distribution is *P* (***s***|***J***, ***h***).

Specifically, we utilize the Gibbs sampling scheme: starting from a random initial state ***s*** = [*s*_1_, *s*_2_, …, *s*_*M*_]^T^, at each step we randomly take an index *k* in the *M* nodes, and update the value of *s*_*k*_ by its conditional distribution given all other nodes *P* (*s*_*k*_ |*s*_1_, …, *s*_*k*−1_, *s*_*k*+1_, …, *s*_*M*_, ***J***, ***h***). After a burn-in period of steps to allow the Markov Chain to equilibrate, the sequence of samples is an empirical distribution of *P* (***s***|***J***, ***h***) and can be used to estimate the gradient Eq. 4, Eq. 5.

In each sampling step, computing the marginal distribution is of complexity *O* (*M*). For accurate cross-correlation estimation, the required number of samples scales as *O* (*M* ^2^), without considering the burn-in period of the MCMC process. Therefore, the overall computational complexity is at least *O* (*M* ^3^). The MCMC method applies to medium-sized networks up to 30 − 50 nodes.

##### Pseudolikelihood method

The major challenge in scaling the inference to large networks is computing the partition function *Z* in the distribution *P* (***s***| ***J***, ***h***), which involves exponentially many terms with the network size *M*. An alternative approximation method called pseudolikelihood was developed originally for spatial statistics and has been adapted to spin network problems [15, 16, 29]. Pseudolikelihood methods approximate the distribution *P* (***s*** |***J***, ***h***) as the product of the conditional distributions of each individual variable, therefore bypassing the need to compute *Z*. The approach provides exact inference in the limit of an infinite number of samples [15].

Specifically, we denote ***s***_*\k*_ = [*s*_1_, …, *s*_*k*−1_, *s*_*k*+1_, …, *s*_*M*_]^T^ as the state ***s*** except *s*_*k*_. The pseudolikelihood function for the inference problem of a single condition is

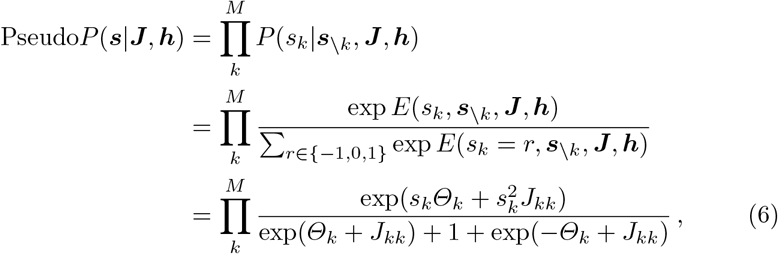

where *Θ*_*k*_ = *h*_*k*_ + ∑_*j*≠*k*_ *J*_*jk*_*s*_*j*_ is the effective field conditioned on all other nodes ***s***_*\k*_. The pseudolikelihood function decouples the mutual dependence between nodes, thus avoiding the computation of the exponentially complex partition function *Z*. As a cost, the pseudolikelihood function in general does not sum to 1, and thus is not a probabilistic distribution; this is where the “pseudo” name comes from. The gradient for the log pseudolikelihood objective function is

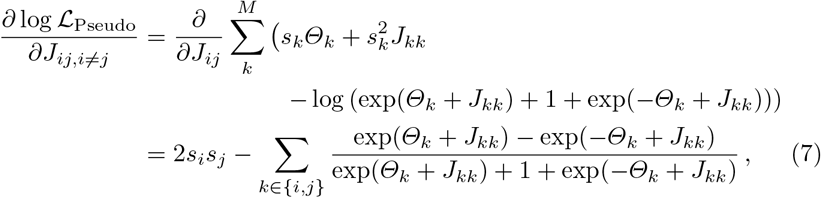

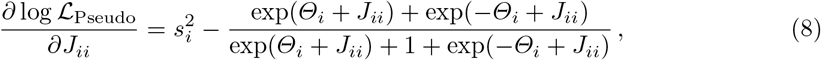

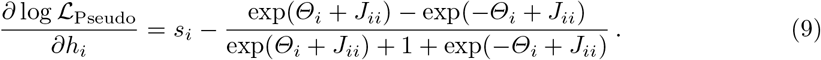

The computational complexity of the gradient computation scales with *O* (*M* ^2^). Also, note that the effective field *Θ*_*k*_ depends on each cell state ***s***, so the gradient computational scales with the total number of cells. Practically, the pseudolikelihood method is highly scalable and even applies to networks of thousands of nodes. The key approximation of pseudolikelihood is to use the empirical distribution to approximate ***s***_*\k*_. The approximation is more accurate when the number of samples is high. Typically, the number of observed samples is far lower than the total number of possible states (3^*M*^). Therefore, exact maximum likelihood and MCMC maximum likelihood are preferable when they are computationally feasible.

#### Assigning directions to the inferred interactions

Identifying directions of interactions from stationary single-cell data has been a difficult problem [30]. The form of pseudolikelihood is closely related to regression models, enabling the assignment of directionality to the inferred network. The distribution of a single node conditioned on all other nodes

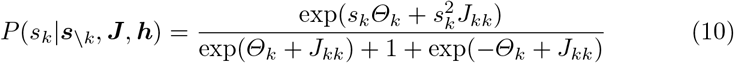

can be interpreted as a regression problem, where we predict the state of the dependent variable *s*_*k*_ using other variables are predictors. The interactions *J*_*ij*_ are the coefficients of the regression problem. If gene A predicts gene B better than B predicts A, this suggests a regulation direction of A to B. To compute the directional network, the gradient estimation Eq. 7 can be simply replaced with an asymmetric version of

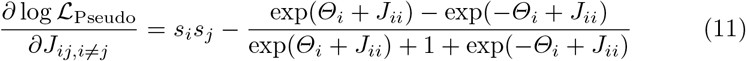

While the algorithm can infer a directed network and achieve high accuracy on synthetic benchmarking datasets [9], it is essential to interpret any directed model from stationary single-cell RNA-seq data with caution. There are several constraints.

First, directed networks cannot define a stationary distribution on transcriptional states when time information is not supplied (*see* **Note 5**). In the context of probabilistic graphical models, directed models are always constrained to be acyclic, i.e., with no cycles. Such a constraint is reasonable in the field of causal inference, where the circularity of causal relations is rare. However, in cellular regulatory networks, feedback loops are prevalent to maintain homeostasis or to achieve signal amplification.

Second, for steady-state snapshot data, an undirected model can adequately explain the observed distribution of transcriptional states (*see* **Note 6**). Identifying directions, in principle, requires the dataset to contain dynamics, even if the time information is not explicitly measured in experiments.

Third, the direction assignment hinges on the key assumption that the asymmetry of predictive power indicates the direction of information flow. The empirical assumption has been adapted in multiple methods such as GENIE3 and GRNBoost2 [30–32], but other factors may also create such asymmetry, for example, the noise level of gene expression. High-expression genes with low observation noise may appear to predict the noisy low-expression genes better compared with the prediction in the reverse way.

#### Identifying gene regulators of programs

The directed network inference algorithm enables associating gene-level networks with program expressions to reveal their mechanisms of regulation. In principle, each gene program can be treated as a “pseudo-gene” and included in the directed network construction. However, the intuitive approach has two major issues. First, as we have discussed, the inferred direction of interaction also depends on the noise level of genes or programs. As a weighted average of multiple genes, the program has a significantly lower noise level than single genes and may spuriously appear as a universal regulator, even though the causal direction should be the reverse. Second, allowing each gene program to have a condition-specific bias term *h*^(*n*)^ would cause the bias term to explain a large proportion of variance with little interpretive value.

To address these issues, we developed a framework where the conditional dependence is computed only in the direction of gene-to-program rather than program-to-gene. Each gene program is also limited to a single global bias *h* that remains fixed across conditions. With these constraints, the model takes on a regression form similar to the pseudolikelihood approach. Specifically, for program expression level *w* ∈ *{*−1, 0, 1*}* and gene expression state ***s***, the log likelihood function of the program state is

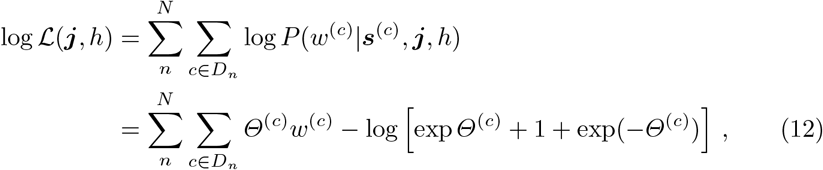

where 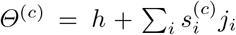is the cell-specific effective field on the program created by single genes and the global bias *h*. The form is similar to Eq. 10, thus the same optimization procedures can be applied.

### 3.2 Identification of gene programs

#### Non-negative matrix factorization

We apply orthogonal non-negative matrix factorization (oNMF) to identify gene programs because its mathematical constraints yield highly accurate, biologically interpretable approximations of single-cell transcriptional states while preventing overlap between programs. Compared to typical matrix factorization methods like principal components analysis (PCA), oNMF imposes two constraints on the gene programs: non-negative weights and orthogonality, both of which are essential for building interpretable interaction models.

1. Linearity and non-negativity: Each program is expressed as a linear combination of individual genes with strictly positive weights, ensuring that it captures a coherent set of co-expressed genes. Enforcing positive weights eliminates the interpretational ambiguity of having negatively weighted genes.
2. Orthogonality: Enforcing orthogonality makes programs non-overlapping, so each can be interpreted as a distinct biological functional module. The profile of each cell is described as a combination of these module activities. In the absence of orthogonality, a cell would be dominated by a single broad program encompassing all genes expressed by that cell type. Within D-SPIN, shared genes between programs would introduce confounding interactions, so the independence conferred by orthogonality is essential.

Formally, given a non-negative gene matrix *X* ∈ ℝ^*C×M*^ with *C* cells and *M* genes, and a chosen program number *K*, oNMF solves the following optimization problem.

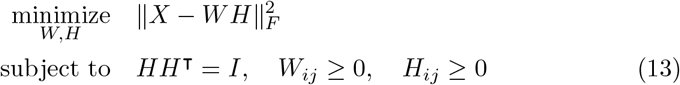

where *H* ∈ ℝ^*K×M*^ is the gene program representation, *W* ∈ ℝ^*C×K*^ is the cell state represented on the gene programs and 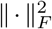is the matrix Frobenius norm.

We implemented a published iterative algorithm [21], with *W* and *H* randomly initialized by uniform distribution on [0, 1]. The random initialization of *H* is orthogonalized using singular value decomposition (SVD) and takes the absolute value before the start of the iteration.

Because the oNMF objective implicitly gives more weights to genes and cells that contribute more to the overall variance, highly abundant cell states and highly expressed genes can dominate the fit. To prevent overfitting to overrepresented cell states, we subsample the single cells to reduce the proportion of highly abundant cell states (*see* **Note 7**). We also normalize the gene expression by the standard deviation of each gene to balance the contribution of highly expressed genes (*see* **Note 8**).

Exact non-negative matrix factorization is an NP-hard problem [33]. Practical algorithms of oNMF rely on heuristic approximations of iterative matrix update, and their solutions can vary with random initialization. To improve robustness, we run oNMF with multiple random seeds and derive a consensus decomposition. For example, we take each row of *H* (feature vector) across repeats of different random seeds, and perform K-means clustering to compute the consensus composition of each gene program. We then define the consensus feature vector as the average of all feature vectors in each cluster, and assign each gene to the consensus program where it has the highest weights.

#### Evaluating gene program expression

Although each oNMF run produces a cell-by-program matrix *W* along with the program composition *H*, after defining the consensus program (which alters *H*), we need to re-evaluate *W* for the consensus program. At this stage, we can also include user-defined programs derived from prior biological knowledge. We then rerun the factorization Eq. 13 with the known gene composition of every program. Mathematically, we impose an additional constraint of knowing which genes belong to each program, i.e., the nonzero entries of *H*.

With known gene assignments, the objective function can be split by each program and optimized individually as

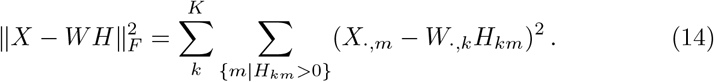

For every program *k*, we only need to consider genes *m* that are known to belong to the program (*H*_*km*_ *>* 0). For notation simplicity, we denote *X*^(*k*)^ as the matrix of genes that are known to belong to program *k, W* ^(*k*)^ as the cell expression level on program *k*, and *H*^(*k*)^ as the weights on genes in the program *k*. The optimization problem on program *k* is

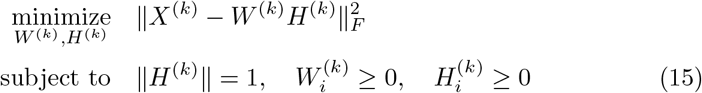

This is essentially a rank-1 non-negative matrix factorization (NMF) problem and has well-established implementations in the Python `scikit-learn` package.

### 3.3 Case study: dissecting the immune dictionary dataset with D-SPIN

#### Overview of the immune dictionary dataset

As a demonstration of D-SPIN, we apply the framework to a public immune dictionary dataset to dissect the action of cytokines on multiple immune cell types *in vivo* [24]. In the experiments, 86 different cytokines were injected into mice, and skin-draining lymph nodes were collected and profiled with single-cell RNA-seq 4 hours after injection. The full dataset is composed of 17 immune cell types with 386,703 single cells. For simplicity, we focus on a subset of data of CD4, CD8, and regulatory T cells and 12 cytokines that were found to induce specific polarization states of all three cell types [24]. The selected cytokines include type I/II interferons (IFN*α*1, IFN*β*, IFN*ϵ*, IFN*κ*, IFN*γ*), IL-1 family (IL-1*α*, IL-1*β*, IL-18, IL-36*α*) and other interleukins (IL-2, IL-12, IL-15), each driving the cell population to distinct polarization states as rendered on the t-distributed stochastic neighbor (t-SNE) embedding (Fig. 2A).

**Fig. 2.**
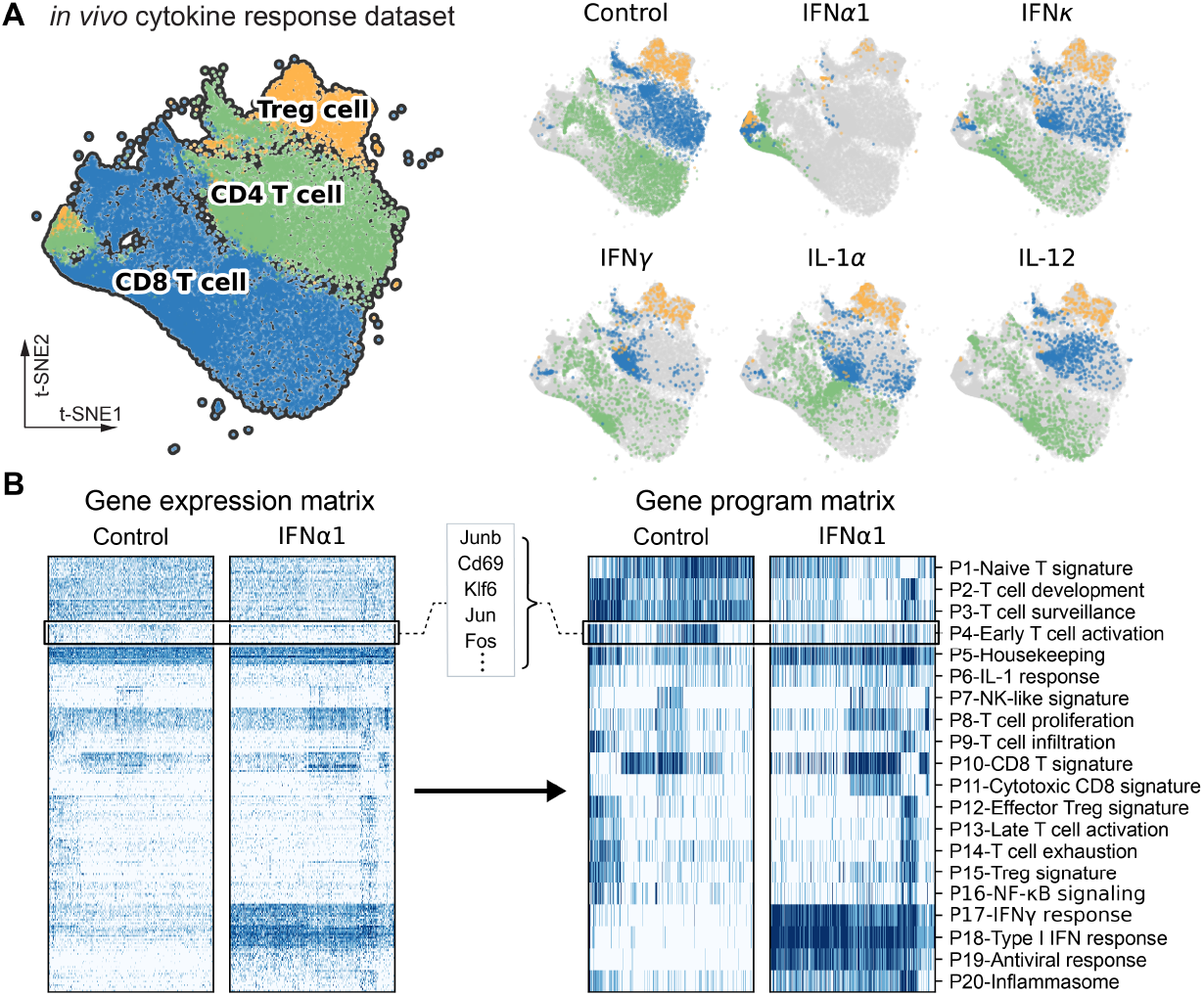
Overview and program discovery for the immune dictionary dataset. (A) t-distributed stochastic neighbor (t-SNE) embedding for a subset of cell populations from the immune dictionary dataset [24]. The subset dataset includes CD4, CD8, and regulatory T cells treated by 12 different cytokines, as well as corresponding control samples treated by phosphate-buffered saline (PBS). (B) Heatmaps of gene expression and discretized gene program levels for control and IFN*α*1-treated samples. The gene programs are weighted averages of single-gene expressions that characterize and denoise the major expression pattern of the gene matrix.

#### Program-level network model partitions cytokine categories

D-SPIN extracted 20 gene expression programs from 2,000 highly variable genes that capture the principal patterns of co-expression (Fig. 2B). These include broad cell-type signature programs composed of known gene markers such as P10-CD8 T signature (Nkg7, Cd8b1, Cd8a) and P15-Treg signature (Il2rb, Il2ra, Foxp3), as well as programs linked to specific cell polarization states and cytokine responses. For example, P18-Type I IFN response program is enriched for interferon-stimulated genes (ISGs) (Ifi203, Sp100, Isg15) that reflect direct response to the interferon stimulation. P6-IL-1 response program contains genes directly induced by IL-1 (Stat3, Bcl3, Cebpb) together with multiple receptors of other cytokines (Ifngr1, Il2r, Il6st, Il4ra), indicating the complex *in vivo* crosstalks between signals and cell types.

The inferred program regulatory network has a distinct modular structure with four network modules of tightly interacting gene programs (Fig. 3A). Modules are defined as subsets of nodes where interactions between nodes in the same module outnumber interactions between different modules. Modules can be identified through iteratively updating partitions of the network to maximize the modularity score, such as the Leiden algorithm (*see* **Note 9**)[34]. Each module aligns with a major cell type in the population. For example, the module associated with regulatory T cells contains programs for Treg, effector Treg, T cell exhaustion, and NF-*κ*B signaling. There are two modules of CD8 T cells; one is standard CD8 T functions, including cytotoxic and NK-like programs; the other is antivirus and interferon responses.

**Fig. 3.**
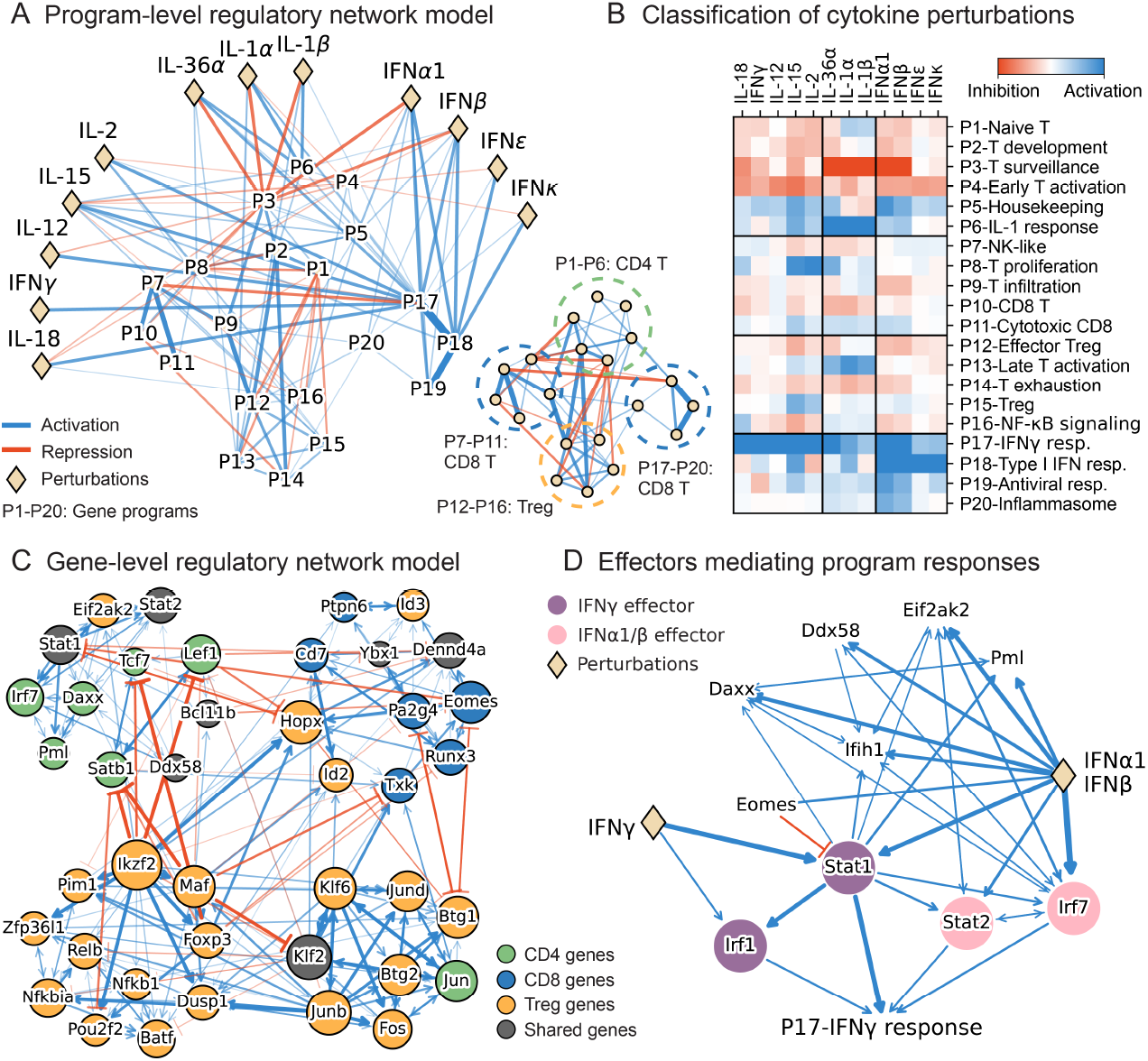
Program-level and gene-level regulatory network inferred by D-SPIN. (A) Diagram of D-SPIN-inferred network model on gene programs. The network is partitioned into 4 modules, each associated with a T cell type in the population. (B) Heatmap of the program response of each cytokine. Clustering of the response partitions the cytokines into 3 major categories. (C) Diagram of the core subnetwork of the D-SPIN-inferred gene network model. The node sizes scale with the number of identified interactions. The network is partitioned into 4 modules, each primarily composed of genes that have elevated expression in one specific T cell type. (D) Interaction diagram for the subnetwork of IFNα1/IFNβ and IFNγ acting on the program P18-IFNγ response. The D-SPIN model shows that Type I and Type II interferons have different effectors to activate the program.

Together with the network, D-SPIN infers the interactions of the 12 cytokines with the regulatory network that drives the observed cell state transitions (Fig. 3B). The cytokines are grouped into three categories by perturbation responses: (i) Th1-polarizing cytokines (IL-2, IL-12, IL-15, IL-18, IFN*γ*); (ii) IL-1 subfamily (IL-1*α*, IL-1*β*, IL-36*α*); and (iii) type I interferon (IFN*α*1, IFN*β*, IFN*ϵ*, IFN*κ*). The classification agrees with the cell polarization analysis of the original immune dictionary study [24]. Cytokines in the same group share dominant perturbation responses, yet retain distinctive secondary signatures. For instance, all four type I interferons strongly activate P18-Type I IFN response program, while IFN*α*1 and IFN*β* additionally induce stronger P17-IFN*γ* response and suppress P3-T cell surveillance. The different responses reflect the potency difference between the two interferon subgroups. IFN*α*1*/β* are universally produced by most cell types and can trigger global antivirus responses when their pattern recognition receptors (PRRs) are stimulated [35]. By contrast, IFN*ϵ/κ* are constitutively expressed at the mucosal and cutaneous barrier and signal with much lower efficacy due to reduced affinity for the IFNAR2 receptor [36].

Collectively, the program-regulatory network constructed by D-SPIN organizes the core cellular functions and cytokine response pathways into an interpretable network diagram. By computing cytokine perturbation responses on this network, the D-SPIN model collapses the cytokine effect on different T cell types into a single vector of coefficients on gene programs. The response vector provides a fingerprint for each cytokine that registers how the perturbation shifts the entire landscape of T cell states. These unified fingerprints of responses provide systematic classifications of cytokine actions and reveal the different signatures of each cytokine associated with their physiological roles.

#### Gene-level network model identifies key effectors of interferon responses

To identify gene-level regulators of cytokine actions and gain a fine-grained view of the regulatory network controlling cytokine signaling, we also construct a gene-level D-SPIN network model. Upon cytokine stimulation, regulation is carried out through both DNA-binding interactions of TFs and signal transduction cascades with kinases and phosphatases. Therefore, we selected 115 highly expressed regulatory genes of 91 TFs, 14 kinases, and 10 phosphatases for the network construction.

Among all these regulatory genes, D-SPIN identifies a core network of 41 genes, highlighting these signaling hubs in the cytokine response (Fig. 3C). The core subnetwork is partitioned into four modules, where two modules are dominated by genes highly expressed in regulatory T cells, and the other two are dominated by CD4 and CD8 T-cell-associated genes. The top signaling hub genes with the highest number of interactions participate in multiple core immune functions, including early signal response by AP-1 (Junb, Jun), cell lineage maintenance (Ikzf2, Hopx), differentiation (Klf2, Klf6), and T cell quiescence (Btg1, Btg2).

Furthermore, D-SPIN identifies regulators for each gene program to couple them to the gene-level regulatory network, enabling the model to nominate the key effectors for the observed program-level responses. As an example, both type I interferon IFN*α*1*/β* and type II interferon IFN*γ* induce the activation of P17-IFN*γ* response in the *in vivo* experiment. The circuit model inferred by D-SPIN reveals the different pathways of action for the two types of interferons (Fig. 3D). The IFN*γ*-driven circuit is straightforward, with only Stat1 and Irf1. Stat1 is the most important TF mediating the transcription response of IFN*γ* [37]. Irf1 is also preferentially induced by IFN*γ* and participates in its signal transduction [37, 38]. In contrast, apart from the core effector Stat1, the effect of IFN*α*1*/β* is mediated by Stat2 and Irf7. Stat2 is a core component of type I interferon signaling by forming the ISGF3 complex [37], and Irf7 is the master regulator in immune response of type I interferon signaling [39]. Moreover, the circuit of IFN*α*1*/β* action contains many secondary regulators, indicating that the response is a consequence of cell-cell communications via additional cytokine secretion.

Overall, the gene-level D-SPIN network extracts the core signaling network, highlights cell-type-specific hub genes, and dissects convergent program responses into cytokine-specific circuits. By associating program response to the gene regulatory network, D-SPIN helps to clarify how type I and type II interferons produce overlapping transcriptional signatures while relying on distinct mechanisms of action, therefore nominating key regulatory nodes for modulation of the T cell cytokine responses.

### 3.4 Building network models with the D-SPIN package

#### Gene-level regulatory network models

Here we present the detailed steps of constructing gene-level regulatory network models with the D-SPIN package. For the input `AnnData` object `adata` prepared as described, the basic steps include

~~~
from dspin.dspin import DSPIN
model = DSPIN(adata, save_path, num_spin=adata.shape[1])
model.network_inference(sample_id_key=‘sample_id’,
    method=‘pseudo_likelihood’, directed=True,
    sample_list=None, perturb_matrix=None,
    prior_network=None, run_with_matlab=False,
    params={‘stepsz’: 0.05, ‘lam_l1_j’: 0.01})
model.response_relative_to_control(sample_id_key=‘sample_id’,
    if_control_key=‘if_control’, batch_key=‘batch’)
~~~

The computed regulatory network model is stored in the model object:

– `model.network`: regulatory network ***J***
– `model.responses`: gene-level responses ***h***^(*n*)^
– `model.relative_responses`: response relative to control. The relative responses are preferably compared with the control sample of each experimental batch.

Remarks regarding the parameters in the gene-level model construction:

– `num_spin` sets the number of network nodes. Therefore, setting num_spin to the number of genes will construct gene-level networks.
– `directed` sets whether to infer a directed network. Only the pseudolikelihood method supports directed network inference.
– `prior_network` (optional) is a binary matrix indicating more likely network edges obtained from prior knowledge, such as TF-motif interactions or chromatin accessibility.
– `sample_list` (optional) is used for restricting the model to a subset of conditions or enforcing a specific order of conditions to match the input of the perturbation matrix.
– `perturb_matrix` (optional) sets the putative effect of each perturbation with shape (sample number) * (gene number). In cases where the perturbations have a known effect, such as Perturb-seq, the matrix is used as a prior to guide the perturbation effect inference.
– `run_with_matlab` blocks inference and saves all relevant data to `save_path` instead. The saved data allows running the MATLAB implementation deposited in GitHub for large-scale datasets.
– `params` is a dictionary of hyperparameters for the optimization. For example, the step size of the gradient ascent, the number of training epochs, and ℓ_1_ or ℓ_2_ regularization of the model parameters.

#### Program-level regulatory network models

The steps for program-level network models are similar, with an extra step of gene program discovery.

~~~
from dspin.dspin import DSPIN
model = DSPIN(adata, save_path, num_spin=20)
model.gene_program_discovery(num_repeat=10, seed=0,
    cluster_key=‘cell_type’)
model.network_inference(sample_id_key=‘sample_id’,
    method=‘mcmc_maximum_likelihood’)
model.response_relative_to_control()
~~~

In addition to the computed network ***J*** and responses ***h***, the program-level D-SPIN model also saves the gene compositions of the consensus gene program discovery under `save_path`.

Remarks regarding the parameters in the program-level model construction

– `num_spin` sets the number of programs. Choosing the optimal number of clusters in dimensional reductions has no universal standard for optimal results. In practice, a heuristic is to set `num_spin` about 5x the number of cell types or clusters in the dataset, but not exceeding 40 programs (*see* **Note 10**).
– `cluster_key` is used to balance proportions of different cell types. The package downsamples overrepresented cell types to avoid overfitting to these states (*see* **Note 7**).

#### Identifying gene regulators of programs

Given a program-network D-SPIN object `model_program` and a gene-level D-SPIN object `model_gene`, the code computes the gene regulators in the gene-level network for each program.

~~~
model_gene.program_regulator_discovery(
    model_program.program_representation,
    sample_id_key=‘sample_id’,
    params={‘stepsz’: 0.02, ‘lam_l1_interaction’: 0.01})
~~~

The `adata` objects in `model_program` and `model_gene` should have the same cell metadata adata.obs, to align the gene expression state and program-expression state of each cell. The computed interactions between genes and programs are:

– `model_gene.program_interactions`: regression coefficients ***j*** between genes and programs.
– `model_gene.program_activities`: the global activity *h* for each program.

#### Module discovery and visualization of network models

With the inferred network, the package also includes functions for network module discovery and visualizations.

~~~
import dspin.plot as dp
G, j_filt = dp.create_undirected_network(model_program.network,
    node_names=spin_name, thres_strength=0.05)
module_list = dp.compute_modules(G, resolution=1, seed=0)
dp.plot_network_heatmap(j_mat_filt, module_list,
spin_name_list=spin_name)
dp.plot_network_diagram(j_mat_filt, module_list, pos=None,
    directed=False, weight_thres=0.15,
    spin_name_list_short=spin_name_short)
dp.plot_response_heatmap(model_program.relative_responses,
    module_list, spin_name_list=spin_name,
    sample_list=model_program.sample_list)
~~~

Modules are identified through Leiden clustering, and resolution controls the granularity of detected modules (*see* **Note 9**). For program-level models, `spin_name_list` are the names of gene programs manually specified or model-identified representative gene names in the program. `spin_name_list_short` are index-like short names in graph rendering. For gene-level models, both name lists will be simply gene names.

### 3.5 Working example of D-SPIN on the immune dictionary dataset

Complete Jupyter notebook to analyze the pre-processed immune dictionary dataset and to visualize the inferred regulatory networks available at Caltech Research Data Repository https://doi.org/10.22002/gaaps-axj72. The repository also contains the preprocessed single-cell data, gene program decomposition, and D-SPIN model objects for reproducibility.

## 4 Notes

1. Rigorously speaking, the model contains an ***h***^(control)^ as the activity of individual nodes without the perturbation. The perturbation modulates the node activity with an extra term ***h***^(perturbation)^. As the energy *E* = − ∑_*ij*_ *J*_*ij*_ *s*_*i*_ *s*_*j*_ − ∑_*i*_ *h*_*i*_*s*_*i*_ is a linear function of ***h***, we combine ***h***^(control)^ and ***h***^(perturbation)^ into notation ***h***^(*n*)^ together for simplicity. Thus, for each perturbation condition, ***h***^(*n*)^ should be compared with the control condition to obtain the relative response under the perturbation condition.
2. We discretize the expression into 3 levels with the form [−1, 0, 1] due to the following considerations
  a. A higher number of levels raises computational complexity and makes the model behave more like a Gaussian distribution with a single probability peak, which cannot capture cell populations with multiple distinct cell types.
  b. A simple on-off discretization makes it difficult to capture different levels of responses or dosage effects.
  c. The [−1, 0, 1] scheme creates interpretable self-interaction *J*_*ii*_. With *J*_*ii*_ *>* 0, the node prefers −1 or 1 state, similar to a bistable switch. With *J*_*ii*_ *<* 0, the node prefers the 0 state, similar to negative feedback.
3. Various methods have been empirically used to partition continuous expressions into discrete levels, such as using percentiles or standard deviations. Discretization with K-means was found to perform well [40] and has a clear interpretation of minimizing the variance inside each expression level category. In practice, gene expression distributions sometimes have long tails, which means a small proportion of cells have very high expressions. Clipping each gene expression at an upper percentile, such as the 95th percentile, will help K-means better partition the distribution.
4. Indistinguishable models refer to suboptimal solutions with nearly identical objective function values but differing in parameters. The existence of suboptimal solutions can be quantified by Fisher information, which characterizes how probability distributions shift under parameter changes. Suboptimal solutions correspond to the linear subspaces with small Fisher information eigenvalues. We found in separate research that different perturbations produce different subspaces with suboptimal solutions, and combining information between perturbation conditions helps eliminate these suboptimal solutions, yielding accurate network inference [28].
5. The acyclic constraint of directed probabilistic graphical models is fundamental, as cycles in the conditional dependence between variables will produce inconsistent distributions. As mentioned in the seminal work of Judea Pearl on causal inference [41], even three cyclically dependent variables “will normally lead to inconsistencies”, and the acyclic constraint can ensure consistency of the distributions. Constructing distributions on even two mutually dependent variables requires nontrivial constraints [42].
6. For any distribution *P* (***x***), we can construct a stochastic differential equation (SDE) whose stationary distribution is *P* (***x***) with only symmetric interaction between variables. Therefore, if the single-cell observations are sampled from a stationary distribution, in principle, an undirected model suffices to explain the data. The sketch of such an SDE construction is as follows. The distribution can be written in the form of an exponential family *P* (***x***) = exp [− 2*V* (***x***)*/σ*^2^] */Z*. The distribution is the steady-state solution of the Fokker-Planck equation

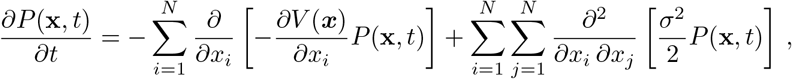

which corresponds to the SDE of

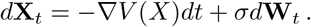 The SDE always has symmetric interactions between variables, as the driving force is the gradient of a potential function.
7. The gene program discovery procedure may overfit to cell states that are over-represented in the cell population. We use two class-balancing strategies from machine learning by subsampling cells based on cluster or cell-type labels.
  a. Equal-sample balancing. We sample an equal number of cells from each cluster to facilitate identifying programs that are expressed only in a small subpopulation.
  b. Square-root balancing. We sample each cluster in proportion to the square root of its size, an empirically effective technique for reducing the proportion of large clusters.
8. Similar to standardizing variables in PCA, we divide the expression of each gene by its standard deviation. Low-expression genes with only a few nonzero entries may become disproportionately large after the normalization. To mitigate this, the standard deviation can be floor-truncated by a certain percentile of the standard deviation of all genes, for example, the 20th percentile.
9. The modularity score in the Leiden algorithm is defined as the density of positive interactions within modules compared to a random network reference [34]. Specifically for networks with both positive and negative interactions, the score function is defined as

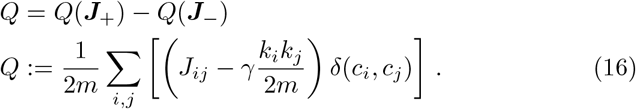 The regulatory network is split into ***J***_+_ for positive and ***J***_−_ for negative interactions by ***J*** = ***J***_+_ − ***J***_−_, where ***J***_+_, ***J***_−_ *>* 0. In the score function *Q, m* represents the sum of all edge weights, *k*_*i*_ denotes the sum of weights for edges connected to node *i, c*_*i*_ is the community to which node *i* belongs, and *γ* is the resolution parameter that controls the granularity of the detected module partitions. Increasing *γ* results in smaller modules containing fewer network nodes.
10. The selection of the number of gene programs is a trade-off between model expressive power, computational complexity, and model interpretability. The optimal choice of program number typically hinges on the specific requirements of the application scenario. Aside from using the number of cell types to roughly estimate the program number, the elbow method also provides a heuristic approach. This method involves plotting a relevant cost function or objective function against the number of gene programs and looking for a point in the plot where the rate of change changes sharply, resembling an “elbow”. For oNMF, the cost function could be the reconstruction error by the gene programs. As the single-cell data is typically noisy, it is recommended to denoise the gene matrix with methods such as data diffusion [43] before computing the reconstruction error for the elbow method.

